# HO-3867 mediated modulation of Extracellular vesicles and Tumor microenvironment: A novel immunotherapeutic strategy for ovarian cancer

**DOI:** 10.1101/2025.05.16.653814

**Authors:** Ganesh Yadagiri, Lakshmi Narasimhan Chakrapani, Shyam Sundaram, Jessica Velasquez, Tiffany Hughes, Gabriela S. Vendrell, Balazs Bognar, Qi-En Wang, David M. O’Malley, David E. Cohn, Karuppaiyah Selvendiran, Kalpana Deepa Priya Dorayappan

## Abstract

**Introduction:** Ovarian cancer (OC) remains the most lethal gynecologic malignancy, with poor long-term survival, largely due to its diagnosis at advanced stages and its high rate of recurrence with resistance to platinum therapies. This study evaluates the therapeutic efficacy of HO-3867, a STAT3 inhibitor, and bevacizumab, an anti-VEGF antibody, alone and in combination in an immunocompetent (IC) syngeneic ovarian cancer progression model.

**Methods:** The ability of HO-3867 to induce macrophage polarization was assessed in RAW264.7 cells, while its impact on anti-tumor immunity was evaluated in splenocytes co-cultured with ID8 and OC ascites cells. Immune modulation, including cytokine induction, extracellular vesicle (EV) release, and endosomal sorting complex required for transport (ESCRT) protein expression, was analyzed *in-vitro* and using *in-vivo* OC model. The therapeutic effects of HO-3867 and bevacizumab were examined in vivo by measuring ascites volume, EV secretion levels, and immune cell populations via flow cytometry.

**Results:** Our results demonstrate that combination therapy significantly reduced ascites accumulation, and limited peritoneal disease spread in ovarian cancer mouse models. EV analysis revealed a decrease in total EVs and CD9+ subpopulations, which correlated with the associated ESCRT proteins involved in EV formation and secretion implicating EVs in tumor progression and immune modulation. Flow cytometry analysis of ascites immune cell populations showed that combination therapy reduced myeloid-derived suppressor cells (MDSCs) and their PD-L1 expression, while enhancing CD8+ T cell cytotoxicity via granzyme B secretion. Cytokine profiling revealed upregulation of IFN-γ and downregulation of IL-6, IL-10 and CXCL-2 suggesting enhanced anti-tumor immune response. Furthermore, HO-3867 promoted macrophage polarization toward tumoricidal M1 phenotype and reduced MDSC expansion *in-vitro*, enhancing anti-tumor immunity.

**Conclusion:** HO-3867 and bevacizumab synergistically reprogram the tumor microenvironment, inhibit EV-mediated progression, and enhance antitumor immunity. These findings support its immunomodulatory potential in ovarian cancer ascites, offering a promising therapeutic strategy.

## Introduction

Ovarian cancer (OC) remains the leading cause of gynecologic cancer-related mortality in the United States, with an estimated 20,890 new cases and over 12,500 deaths in 2025^1,2^. Despite advancements in understanding OC etiology and pathobiology, overall survival remains poor. Key factors contributing to this high mortality include early peritoneal metastasis, high recurrence rates, and the development of chemoresistance. Notably, over 80% of OC cases are diagnosed at an advanced stage with a 5 year overall survival of approximately 50%^3-6^, underscoring the urgent need for improved therapeutic strategies. Identifying key signaling proteins and extracellular vesicles (EVs), such as exosomes, involved in early peritoneal spread^7,8^ could provide critical insights into tumor progression and reveal novel therapeutic targets to enhance treatment efficacy.

Ascites represents a complex tumor microenvironment (TME) that drives ovarian cancer progression and metastasis through immune modulation, contributing to immune evasion and suppression^9^. Recent studies demonstrated an increase in immunoregulatory cytokines, such as IL-6, IL-10, and CXCL-2, which impair immune function and further facilitate tumor immune evasion in the ascites TME^10^. A hallmark of this TME is the accumulation of immunosuppressive myeloid-derived suppressor cells (MDSCs), which co-express myeloid markers (CD11b and Gr-1) and are closely associated with tumor-associated macrophages (TAMs) that express high levels of PD-L1, ultimately reducing CD8+ T cell cytotoxicity through immune checkpoint signaling^11^.

Our research demonstrates that targeting the ascites TME with the small molecule inhibitor HO-3867 in combination with anti-VEGF antibody bevacizumab, effectively reduces tumor progression. HO-3867 significantly decreases extracellular vesicle (EV) secretion, thereby making the TME more susceptible to immune attack by boosting T cell cytotoxicity in syngeneic murine ovarian cancer models. These findings underscore the potential of using HO-3867 and bevacizumab to block EV secretion, reduce immune suppression, enhance anti-tumor immunity, and slow disease progression, providing promising strategies to improve therapeutic efficacy in ovarian cancer. However, the complexity and heterogeneity of the tumor microenvironment (TME) remain significant challenges in developing effective cancer treatments. This study shows that HO-3867 and bevacizumab synergistically reprogram the immunosuppressive ovarian tumor microenvironment, inhibit EV-mediated tumor progression, and enhance antitumor immunity by suppressing MDSC function in ovarian cancer

## Materials and Methods

### Cell lines and culture conditions

The murine ID8 and patient derived ascites R127 ovarian cancer cell lines were cultured in Roswell Park Memorial Institute (RPMI)-1640 medium (RPMI-1640; Gibco) supplemented with 10% fetal bovine serum (FBS), 1% penicillin-streptomycin (Gibco), 1% sodium pyruvate (Gibco) and maintained at 37°C in a humidified incubator with 5% CO_2_ atmosphere. RAW264.7 macrophage cells were cultured under standard conditions in Dulbecco’s Modified Eagle Medium (DMEM) supplemented with 10% FBS and antibiotics.

### Splenocyte isolation and culture methods

Single-cell suspensions of mouse primary splenocytes were prepared by mechanically dissociating spleens from naive female C57BL/6 mice using a sterile syringe plunger in cold complete RPMI-1640 medium, filtering through a 70µM sterile cell strainer into a 50 ml centrifuge tube. The splenocytes were isolated by centrifugation (300 x g for 5 minutes at 4°C), and red blood cells were lysed using ACK lysis buffer (Thermo Fisher) for 3–5 minutes at room temperature. After two washes with complete RPMI-1640 medium, the cells were resuspended in fresh medium, and cell viability was assessed via trypan blue exclusion. The splenocytes were seeded in 24-well plates for co-culture experiments with ID8 cells (tumor and ascites derived) or R127 cells (1:1 ratio) to assess MDSC expansion and immune cell profiling in the presence or absence of HO-3867. All samples were analyzed in triplicates.

### Animal model and tumor induction

Female C57BL/6 mice (6–8 weeks old) were purchased from the Jackson Laboratory and housed under pathogen-free conditions at The Ohio State University (OSU). All animal experiments were approved by the Institutional Animal Care and Use Committee (IACUC) of OSU. Mice were intraperitoneally injected with 2 × 10^6^ ID8 cells suspended in 200 µL of PBS. After 10 days, mice were randomized into four groups (n = 5–7 per group): (1) untreated control, (2) HO-3867-treated, (3) bevacizumab-treated, and (4) HO-3867 + bevacizumab combination therapy. HO-3867 was administered orally (*p*.*o*.) (2.5mg/kg) thrice per week, and bevacizumab was given intraperitoneally (*i*.*p*.) at (5 mg/kg) weekly for 4 weeks.

### Ascites collection and extracellular vesicle (EV) isolation

At the study endpoint, mice were sacrificed, and peritoneal ascites fluid was collected, and total volume was measured in each treatment group. Extracellular vesicles (EVs) were isolated from the ascites using Izon qEV size exclusion chromatography (SEC) columns (Izon Science) according to the manufacturer’s protocol. The isolated EVs were quantified and characterized using Image Stream flow cytometry with Exo-FITC and CD9-PE antibodies (BioLegend) as described previously (REF).

### Transmission Electron Microscopy (TEM)

The metastatic ovarian tumor tissues were processed for TEM imaging as described previously^8^.

### Flow cytometry analysis of immune cells

Splenocytes collected at the end of experimental time points after *in-vitro* co-culture studies were stained for surface markers (CD45, CD11b, Gr-1, PD-L1) and analyzed by flow cytometry. Likewise, Single-cell suspensions from mice ascites collected from different treatment groups were stained with fluorochrome-conjugated antibodies targeting CD11b, Gr-1 (for MDSCs), PD-L1, CD3, CD8, granzyme B, for immune cell profiling. Flow cytometry was performed on a Attune NxT (ThermoScientific) instrument, and data were analyzed using FlowJo software^11^.

### *In vitro* assessment of Macrophage polarization

RAW264.7 cells were treated with HO-3867 (2.5–5.0 μM) for 24 hours and stained for CD11b, F4/80, and CD86 antibodies to assess the induction of M1 macrophage polarization after treatment by flow cytometer (Attune NxT ThermoScientific), and data were analyzed using FlowJo software^11^.

### Annexin V/PI apoptosis assay

Primary patient derived R127 ascites cells were treated with HO-3867 (5-10 μM) for 24 hours. Apoptosis was evaluated by Annexin V-FITC and propidium iodide (PI) staining according to the manufacturer’s instructions using flowcytometry.

### Cytokine and ESCRT protein analysis ascites and R127 cells

Immune modulatory cytokines, IL-6, IFN-gamma, CXCL-2, IL-10, were analyzed by ELISA in cell culture supernatants and mice ascites collected at the end of the treatment period. Samples were assessed in triplicates and analyzed for significant differences using the one-way ANOVA. Likewise, ESCRT machinery proteins associated with EV-formation and trafficking (HRS, VSP4A, VSP28, VSP36 and CHMP4B were analyzed in cell lysates collected from different treatment groups both *in-vitro* and *in-vivo*.

### Statistical analysis

All data were expressed as mean ± standard deviation (SD). Statistical significance was determined using one-way ANOVA followed by Tukey’s post hoc test, where appropriate. p-values < 0.05 were considered statistically significant. GraphPad Prism software (version 10) was used for all analyses.

## Results

### Effect of HO-3867 and bevacizumab treatment in tumor progression in syngeneic immunocompetent mouse model

In ovarian cancer, ascites is linked to tumor progression with cancer spread in the peritoneal cavity leading to aggressive tumor progression and treatment resistance^12^. HO-3867 is a synthetic curcumin analog that inhibits STAT3 in ovarian cancer, while bevacizumab, a monoclonal antibody targeting VEGF, is used in combination with chemotherapy in advanced or recurrent disease to improve drug uptake by lowering interstitial fluid pressure. In our current study, we evaluated the therapeutic efficacy of HO-3867 and bevacizumab alone and in combination, to assess their role in limiting peritoneal spread of disease in an ovarian cancer progression model (Fig.1 A). Metastatic mesenteric tumor nodules were found to be decreased in treatments compared to control as shown in Fig. 1B. The mean volume of ascites accumulation in the combination treatment group(1.4±1.08ml) was significantly reduced when compared to monotherapy (HO-3867-2.8±1.48ml and bevacizumab-5±3.31ml) treatment and untreated control group(13.2±1.30ml) animals suggesting that the combination HO-3867 and bevacizumab acts synergistically in regulating the spread of the disease in the peritoneal cavity as depicted in **Fig. 1 C**

**Figure 1.**
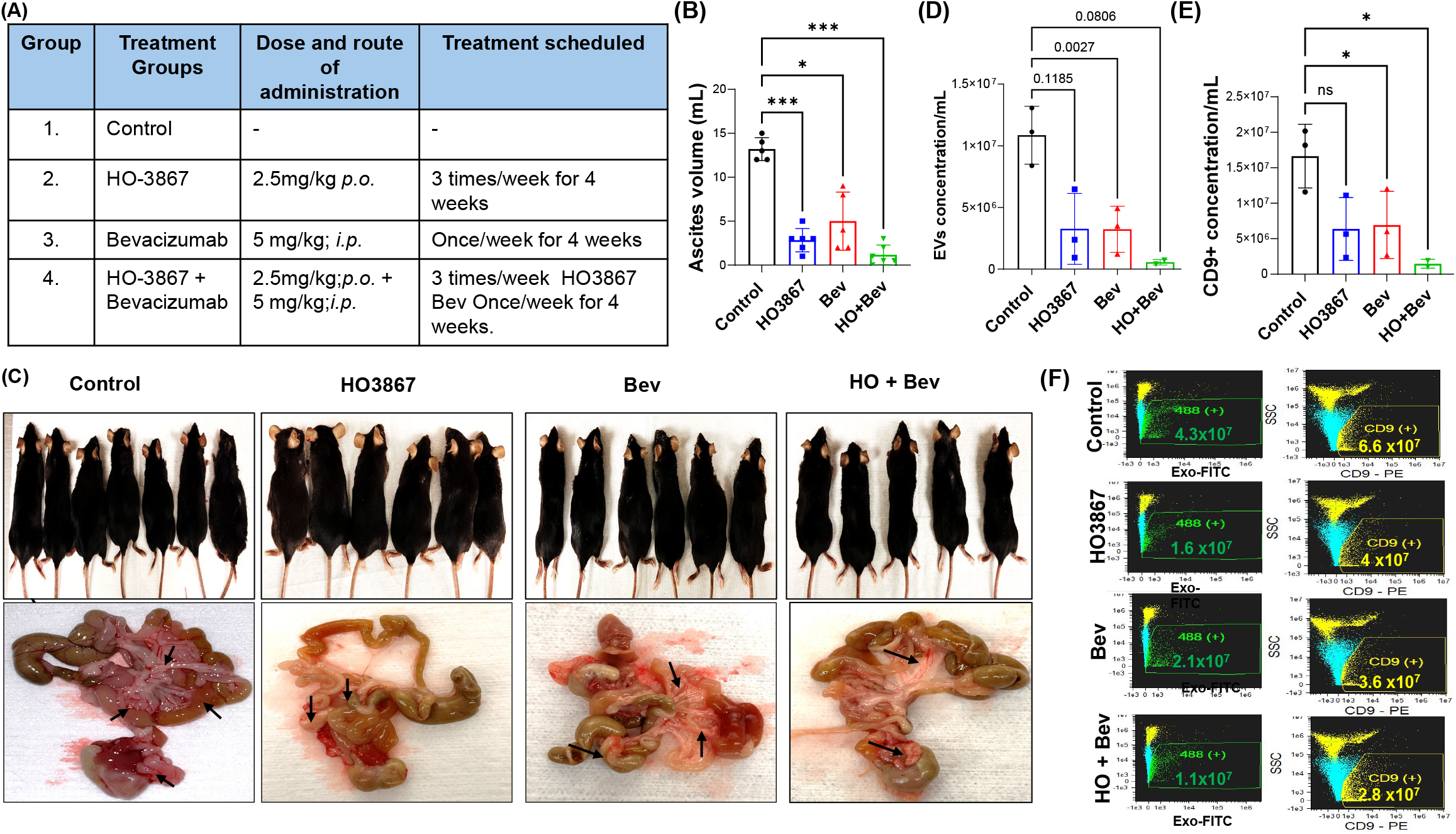
Assessment of therapeutic efficacy of Treatment strategy and EV characterization in ID8 ovarian cancer bearing immunocompetent mice. **A)** Schematic table showing the treatment groups: control, HO-3867, bevacizumab and the combination of HO-3867 + bevacizumab. **B)** Quantification of ascites volume at the experimental endpoint. **C)** Representative images of tumor bearing mice showing differences in abdominal distention and mesenteric nodules across the treatment groups. **D)** Concentration of EVs isolated from ascites, quantified by Image stream flow cytometry analysis. E). CD9+ EV population **F)** Representative image stream flow cytometry showing Exo-FITC+ populations from different treatment groups. Data represented by mean ± SD of 5-6 mice per group and statistical analyses between the groups were done by one way analysis of variance (ANOVA) followed by Tukey post-hoc test. \**P <* 0.05 (Control vs HO-3867, bevacizumab and combination of HO-3867 and bevacizumab).

### Extracellular vesicles (EVs) characterization from mice ascites

Image stream flowcytometry analysis on EVs isolated by qEV-Izon SEC columns from control, HO-3867, bevacizumab and combination treated mice ascites showed significant differences in the total concentration of EVs released in the TME in the combination treatment group when compared to monotherapy treatment and untreated control groups. Interestingly, we found a suppression of total EVs and the EV CD9+ve sub-population in the combination treatment group indicating that CD9+ve exosomes are involved in EV cargo transfer in the ovarian cancer ascites TME as shown in **Fig1D-F**.

### Cytokine modulation in ovarian cancer ascites tumor microenvironment

Ovarian cancer ascites contains cytokines and growth factors from the tumor which accumulate and create an immunosuppressive environment^9,11^. TEM analysis of the metastatic tissues; diaphragm and mesentery reveal a drastic reduction in the vesicle formation on treatment with HO-3867 as compared with untreated control tissues (Fig. 2A) suggesting HO-3867 to be effective in regulating the ESCRT machinery proteins though further studies are warranted to its specific inhibitory role of the ESCRT proteins in EV regulation pathway. Understanding the soluble factors involved in the ovarian cancer tumor environment is key to evaluating cancer progression. In our current study, we aimed to determine the levels of IL6, IFN-γ, CXCL2 and IL10 that could regulate immune suppressive TME. Our results revealed that the combination of HO-3867 and bevacizumab significantly decreased the expression of IL6, IL10 and CXCL2 and increased the expression of IFN-γ in ascites fluid compared to untreated controls, which indicates enhanced anti-tumor immunity by differential modulation of cytokines in treatment groups versus the control group. (**Fig. 2 B**).

**Figure 2.**
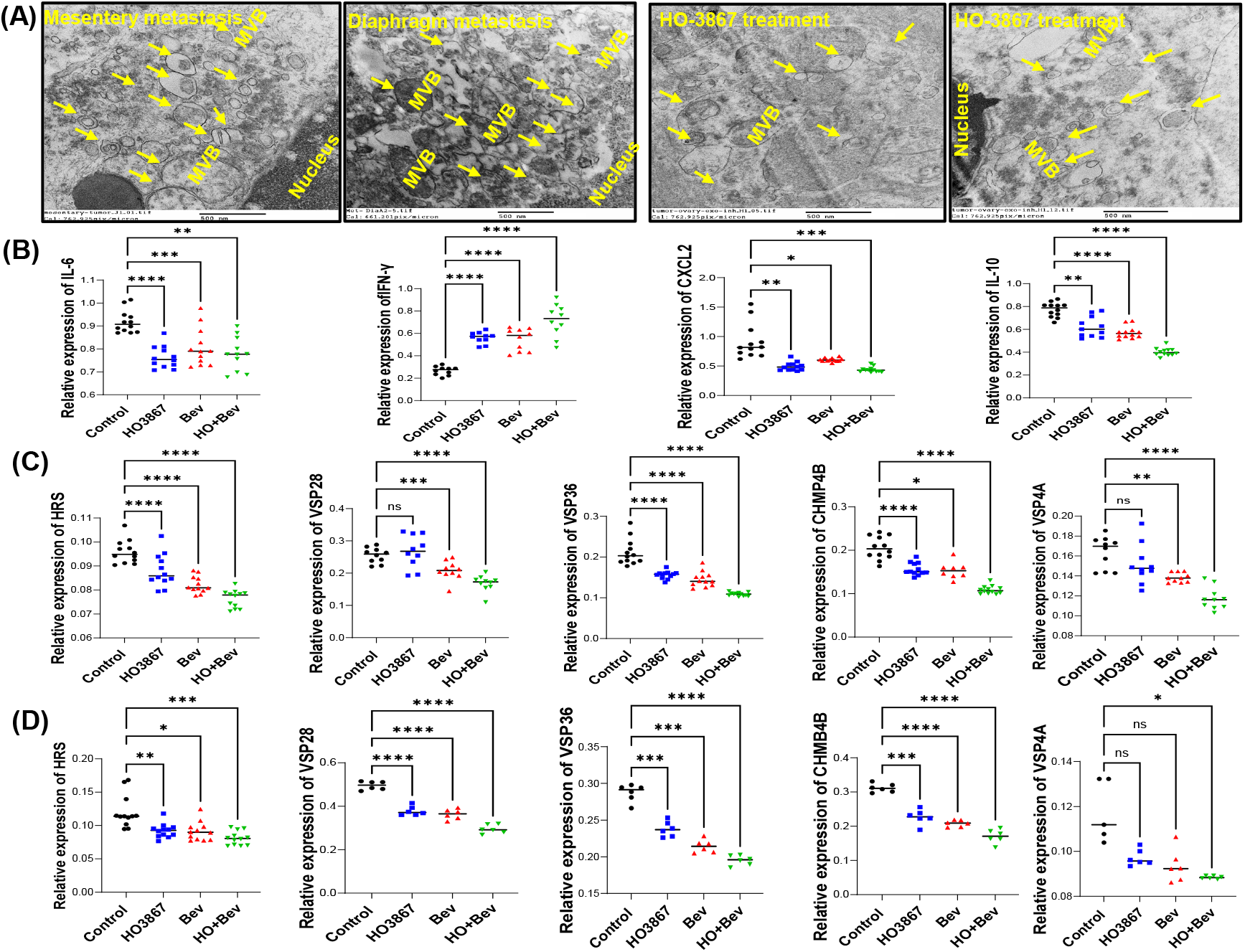
Influence of HO-3867 in ESCRT machinery proteins and soluble factors associated in EV secretion and tumor progression: **A)** TEM images show increased vesicle formation and multivesicular bodies (MVB) in diaphragm and mesentery metastasis of OC progression as compared to the HO-3867 treatment groups (Yellow arrows, scale bar-500 nm. TEM was performed on three independent tumor tissues). **B)** Cytokine analysis in ascites fluid. **C &D)** ESCRT Biomarker protein expression in ascites and R127 cells. ELISA data represented by mean ± S.D of 5 mice and statistical analysis between the groups were done by one way analysis of variance (ANOVA) followed by Tukey post-hoc test. \*\*\**P <* 0.1 (Control vs HO-3867, bevacizumab and combination of HO-3867 and bevacizumab)

### Identifying ESCRT machinery proteins in pre-clinical mice model of ovarian cancer progression

The acellular fraction of ovarian cancer ascites contains EVs and other tumor-promoting factors that could contribute to tumor progression and chemoresistance through immune evasion^13-15^. We previously observed increased EV levels in ovarian cancer ascites and aimed to investigate the EV trafficking proteins HRS (ESCRT-0), VSP28 (ESCRT-1), VSP36 (ESCRT-II) CHMP4B (ESCRT-III), and VSP4A, which constitute different components of the endosomal sorting complex machinery proteins required for transport (ESCRT). Our results demonstrate increased expression of ESCRT proteins in ID8 control ascites, indicating active engagement of the ESCRT machinery proteins in EV formation and secretion in the ascites tumor microenvironment (TME) leading to elevated EV secretion. TME. However, combination treatment with HO-3867 and bevacizumab significantly reduced the expression of these proteins in ascites and in *in-vitro* studies done with patient-derived ascites cells, R127, suggesting a decrease in EV secretion compared to untreated controls (**Fig. 2C &D**). These findings indicate that EVs in ovarian cancer ascites play a role in tumor progression and could be targeted therapeutically to reduce disease burden and improve survival.

### Identification of immune cell population in ascites tumor microenvironment in syngeneic mice model

Ovarian cancer ascites contains both tumor and non-tumor cells, creating a microenvironment that influences tumor cell properties. We aimed to evaluate ascites immune cell components and their role in ovarian tumor progression. Flow cytometry analysis revealed a significant increase in the MDSC population in untreated ID8 ovarian cancer-bearing mice, indicating a highly immunosuppressive TME in ovarian cancer (OC) ascites. Further, combination treatment significantly reduced the MDSC population in ascites (**Fig. 3A** & **B**). Additionally, PD-L1 expression on MDSCs was elevated in the ascites of untreated mice, but treatment with monotherapy HO-3867, bevacizumab, and their combination significantly reduced PD-L1 expression on MDSCs, with the combination showing the greatest reduction. (**Fig. 3C** & **D**). Overall, PDL-1 expression was upregulated in the ascites microenvironment of untreated ID8 cancer bearing mice, contributing to immune evasion and T cell exhaustion. Monotherapy HO-3867 and bevacizumab treatment reduced PD-L1 expression, with combination therapy further downregulating it, suggesting effective reduction of the immunosuppressive tumor microenvironment in ovarian cancer ascites.(**Fig. 3E** & **F**)‥ Additionally, granzyme B released by CD8+ T cells, a measure of cytotoxic function, was reduced in untreated mice, but combination therapy significantly increased granzyme B+CD8+ T cells in ascites, enhancing T cell cytotoxicity **(Fig.3G(to be included in figure**).

**Figure 3.**
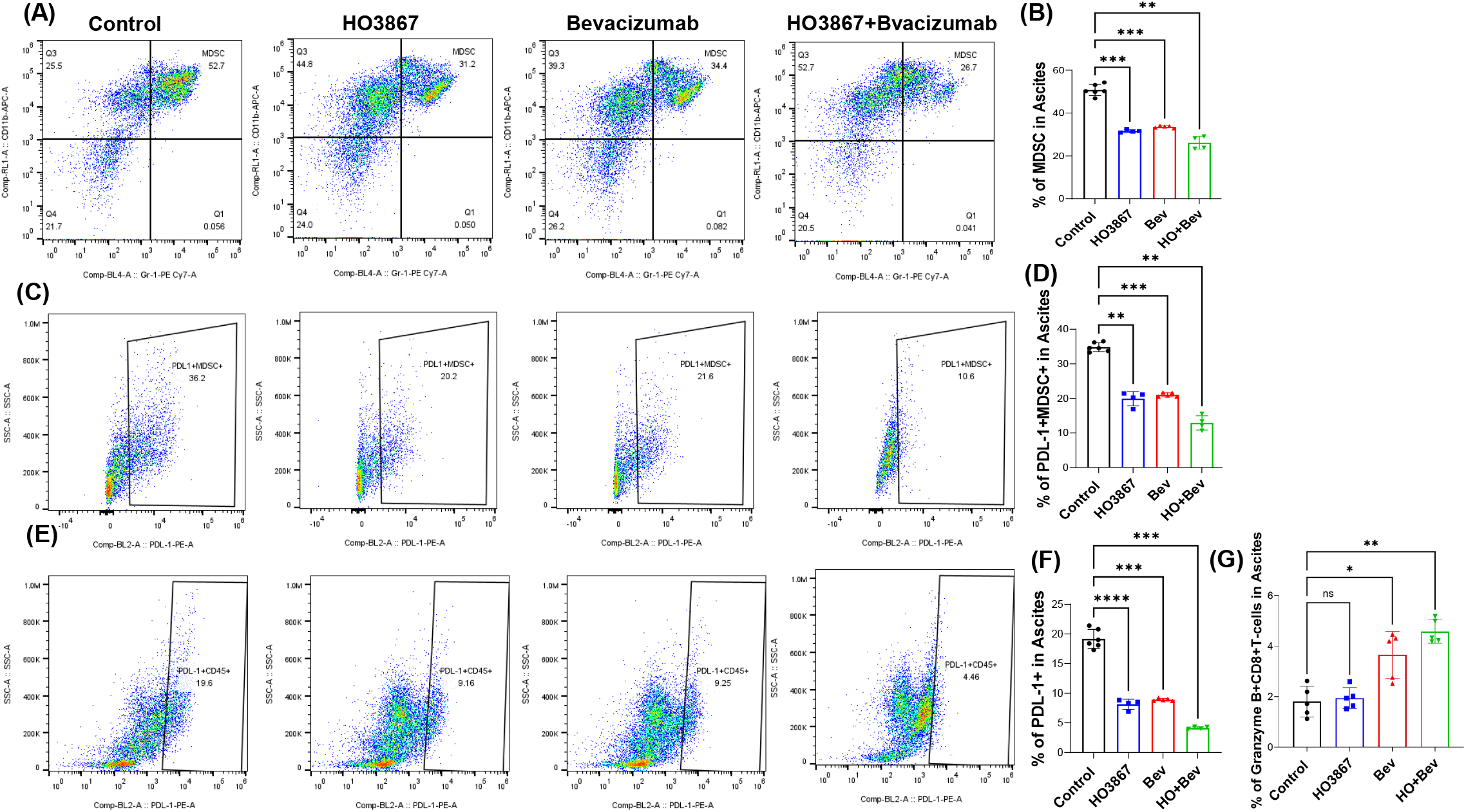
Immunomodulatory properties of HO-3867 and bevacizumab combination in OC progression model in mice. Representative flow cytometry plots showing MDSC (CD11b+Gr-1+) from different treatment groups: **A&B)** Representative flowcytometry plots and quantification of MDSC frequency in ascites across the treatment groups (n=5-6/group). **C&D)** Representative flow cytometry plots and quantification of PD-L1 expression on MDSC (PD-L1+CD11b+Gr-1+) from different treatment groups. **E&F)** Representative flow cytometry plots and quantification of overall PD-L1 expression across cell populations in ascites. **G)** Assessment of Granzyme-B+ on cytotoxic T cells in ascites across the treatment groups(n=5-6/group).

### HO-3867 augments anti-tumor immunity and promotes apoptosis in ovarian cancer cells

The soluble factors in ovarian cancer ascites TME could significantly impact macrophage polarization and antitumor immunity^15,16^. We investigated macrophage polarization effects on RAW264.7 macrophage cells treated with HO-3867 (2.5-5 μM) for the expression of CD11b+F4-80+CD86+ cells by flow cytometry. Our results revealed that HO-3867 significantly increased the macrophage polarization toward M1 phenotype in a dose dependent manner, which might enhance the anti-tumor immune response by reprogramming the tumor progressive M2 phenotype to the M1 tumoricidal phenotype (**Fig. 4A** & **B**). Therefore, manipulating macrophage polarization within the TME can significantly influence antitumor immunity, making it a potential target for cancer therapies. Further, the impact of HO-3867 on MDSC expansion was evaluated in murine splenocytes co-cultured with either ID8 cells or primary ascites cells derived from ID8 in the presence and absence of HO-3867 (5-10 μM) treatment. Our results show that the MDSC (CD11b+Gr-1+) population on cultured splenocytes were significantly decreased in HO-3867 monotherapy treatment in a dose dependent manner (**Fig. 4C** & **D**)‥ Further, analysis of PDL-1 within the MDSC population and total PDL-1 in splenocytes showed remarkable decrease correlating to HO-3867 dose (**Fig. 4E**). These results suggest that HO-3867 may induce anti-tumor immunity by diminishing the presence of immunosuppressive MDSC in the TME. Additionally, we observed a significant increase in apoptosis on R127 ovarian cancer cells co-cultured with splenocytes and treated with HO-3867 when compared with the untreated control group. These findings indicate that HO-3867 induces apoptosis in R127 cells, potentially contributing to its anti-tumor effects (**Fig. 4F** & **G**).

**Figure 4.**
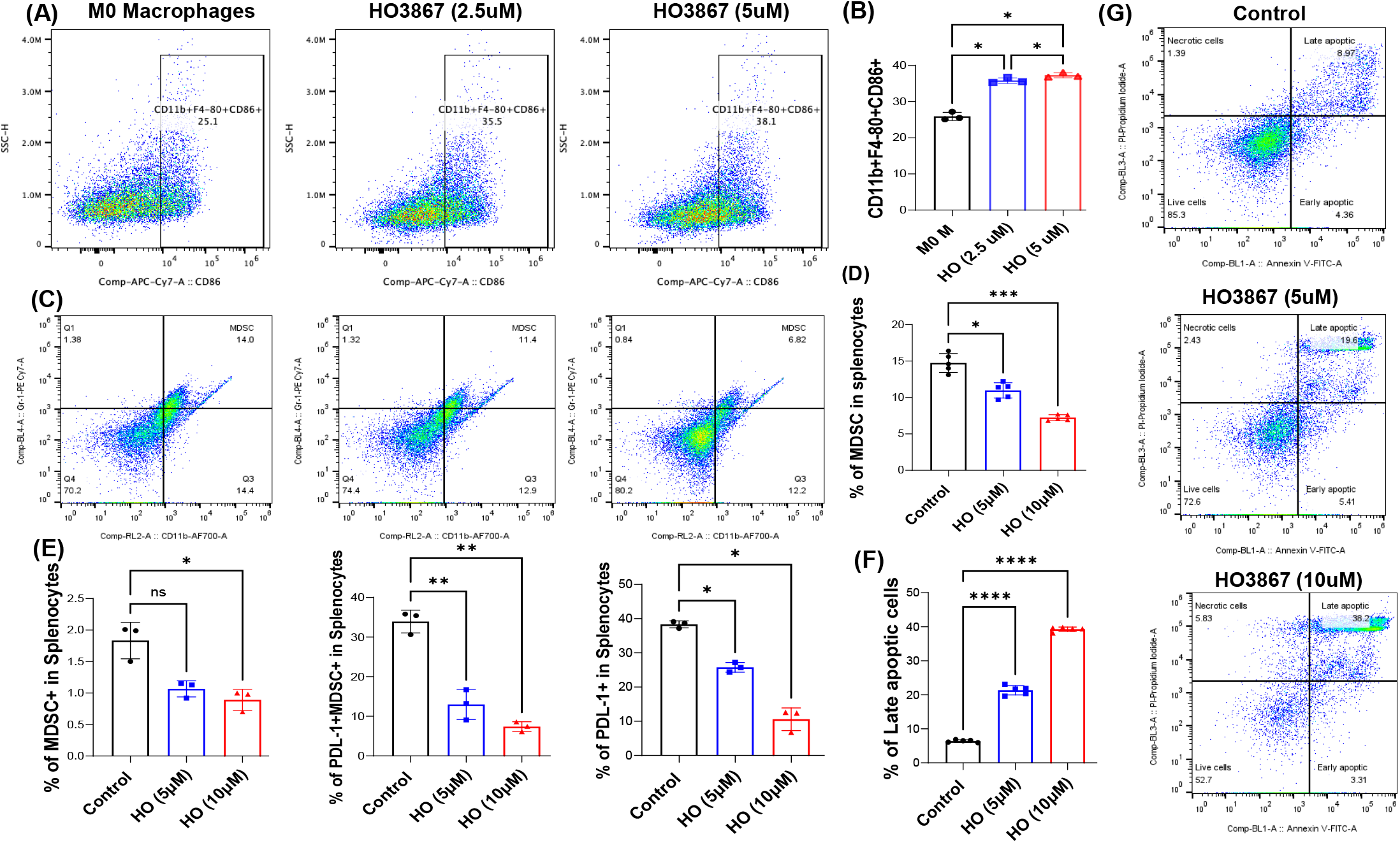
Immunomodulatory properties of HO-3867 *in-vitro*. **A&B)** Representative flow cytometry plots and quantification of M1 macrophage (CD11b+F4-80+CD86+) polarization induction from different treatment concentrations of HO-3867. **C&D)** Representative flow cytometry plots and quantification of MDSC (CD11b+Gr-1+) population reduction from different treatment concentrations of HO-3867 in ID8 primary ascites and splenocyte co-culture experiments. **E)** % of MDSC, PD-L1+MDSC+, and PD-L1+ expression in splenocytes-ID8 cell co-culture from different treatment concentrations of HO-3867. **F&G)** Primary patient derived R127 ascites cells were treated with HO-3867 for 24 hours and evaluated for apoptosis was evaluated by Annexin V-FITC and propidium iodide (PI) staining.

## Discussion

In this study, we present several novel findings on the effects of the DAP compound HO-3867 in combination with bevacizumab in ovarian cancer. Specifically, we demonstrate that (i) the combination of HO-3867 and bevacizumab synergistically reduces EV secretion and ascites fluid accumulation in the peritoneal cavity; (ii) combination HO-3867 and bevacizumab therapy alters cytokine levels and immune signaling proteins in ovarian cancer ascites cells; and (iii) HO-3867 promotes macrophage polarization toward the M1 phenotype while enhancing anti-tumor T cell immunity by reducing the presence of immunosuppressive MDSCs in the TME.

The presence of MDSCs and TAMs has been strongly correlated with poor survival in multiple cancer types, including ovarian cancer^17-19^. These immunosuppressive cell populations contribute to ovarian cancer cell proliferation, angiogenesis, metastasis and drug resistance by producing various oncogenic factors, such as VEGF, cytokines, and CXC-chemokines^20,21^. Our *in-vitro* studies utilizing murine primary splenocytes and ID8 ovarian cancer cells revealed that EVs play a crucial role in establishing an immunosuppressive TME by promoting MDSC expansion and impairing T cell cytotoxicity, thereby facilitating immune evasion^22^. Previous studies from our group have demonstrated that the DAP compound HO-3867 effectively suppresses tumor growth in ovarian and endometrial cancers^23-25^. In this study, we further investigated the anticancer efficacy of HO-3867 in combination with bevacizumab, focusing on its immunomodulatory effects in clinically relevant ovarian cancer mouse models.

Emerging evidence suggests that TAMs represent a significant component of the ovarian cancer ascites microenvironment and play a pivotal role in tumor progression and metastasis^26,27^. Recent preclinical and clinical studies indicate that targeting TAMs and their polarization status may offer a promising strategy for enhancing ovarian cancer treatment^28-31^. Our current findings demonstrate that HO-3867 significantly promotes macrophage polarization toward the M1 phenotype, which may enhance the anti-tumor immune response by reprogramming tumor-promoting M2 macrophages into tumoricidal M1 macrophages, thereby improving treatment efficacy.

In summary, our study highlights the role of ovarian cancer ascites in modulating immune cell function through EV secretion by affecting the ESCRT machinery proteins. Targeting the ESCRT pathway to regulate extracellular vesicle (EV) release holds promise in cancer therapy, where EVs contribute to tumor progression, metastasis, and therapy resistance by transferring oncogenic proteins and other bioactive molecules. By inhibiting ESCRT components we show that the release of tumor-derived EVs can be reduced, potentially disrupting harmful cell-to-cell signaling within the tumor microenvironment. This provides evidence that the STAT3 inhibitor HO-3867, in combination with bevacizumab, synergistically reprograms the immunosuppressive TME, inhibits EV-mediated tumor progression, and enhances anti-tumor immunity. These findings offer preclinical support for the immunomodulatory effects of this combination therapy in ovarian cancer and underscores an innovative therapeutic strategy with the potential to improve patient outcomes.

